# Multiplexed Methylated DNA Immunoprecipitation Sequencing (Mx-MeDIP-Seq) to Study DNA Methylation Using Low Amounts of DNA

**DOI:** 10.1101/2023.01.09.523163

**Authors:** Inam Ridha, Chenxi Xu, Yunro Chung, Jin G Park, Joshua LaBaer, Vel Murugan

**Affiliations:** School of Biological and Health Systems Engineering, Arizona State University, Tempe, AZ, United States; Center for Personalized Diagnostics, Biodesign Institute, Arizona State University, Tempe, AZ, United States; College of Health Solutions, Arizona State University, Phoenix, AZ, United State

## Abstract

DNA methylation is an epigenetic mark that has a crucial role in regulating gene expression. Aberrant DNA methylation results in severe diseases in humans, such as cancer, autoimmune disease, atherosclerosis, and cardiovascular diseases. Whole-genome bisulfite sequencing and methylated DNA immunoprecipitation are available to study DNA methylation changes, but they are typically used on a few samples at a time. Here, we developed a novel method called Multiplexed Methylated DNA Immunoprecipitation Sequencing (Mx-MeDIP-Seq), that can be used to analyze many DNA samples in parallel, requiring only small amounts of input DNA. In this method, 10 different DNA samples were fragmented, purified, barcoded, and pooled prior to immunoprecipitation. In a head-to-head comparison, we observed 99% correlation between MeDIP-Seq performed individually or combined as Mx-MeDIP-Seq. Moreover, multiplexed MeDIP led to more than 95% normalized percent recovery and a 25-fold enrichment ratio by qPCR, like the enrichment of the conventional method. This technique was successfully performed with as little as 25 ng of DNA, equivalent to 3400 to 6200 cells. Up to 10 different samples were processed simultaneously in a single run. Overall, the Mx-MeDIP-Seq method is cost-effective with faster processing to analyze DNA methylome, making this technique more suitable for high-throughput DNA methylome analysis.

## Introduction

Epigenetics studies heritable phenotypic changes, including gene expression, that do not involve changes to DNA sequences [1, 2]. DNA methylation (DNAm) is one of the most widely studied epigenetic marks, as it plays a crucial role in gene silencing, cell fate decisions, and disease development [2, 3]. It involves covalent modification of the pyrimidine ring of cytosine nucleotide at the C-5 position through the addition of a methyl group. DNAm is mainly found as symmetrical CpG dinucleotides in mammalian cells [4, 5]. It is established during gametogenesis and embryo development by recruiting de novo methyltransferase enzymes preserved by maintenance methyltransferases [6, 7]. Changes in DNA methylation continue during adulthood by the recruitment of methyl-CpG binding domain family of proteins, like methyl-CpG binding domain proteins such as MeCP2 in concert with histone deacetylases [8-10]. This epigenetic modification is essential for the normal function of cellular processes such as differentiation and preservation of genomic stability [11, 12]. Aberrant DNA methylation may cause cancer and other diseases such as autoimmune disorders [13, 14] or Alzheimer’s disease [15, 16]. Up-regulation or down-regulation of the methyltransferase enzymes [17], nutrient intake [18, 19], environmental stress such as air pollution, infectious agents, and toxic elements [20-22] are considered among the factors that can cause DNA methylation alteration.

Several techniques study changes in methylation patterns, [23] transforming that information into quantitative and measurable signals [24, 25]. This includes methods to investigate methylation in detail at the level of a single base, such as whole-genome bisulfite sequencing (WGBS) or enrichment of methylated DNA fragments through immunoprecipitations using antibody or methyl binding domain proteins (MBD) [26]. WGBS combines sodium bisulfite conversion of input DNA and high-throughput DNA sequencing [27] to create a comprehensive measurement of all methyl additions to the DNA. Although bisulfite sequencing leads to DNA methylation analysis at single-base resolution [28], the treatment causes substantial DNA degradation and needs excessive purification to remove the sodium bisulfite, thus requiring at least one microgram of input DNA to start the process [29]. Other limitations of WGBS are the high cost of the technology, especially on large sample sizes [30], and the inability of this method to distinguish between 5-methylcytosine (5mC) and hydroxy-methylcytosine (5hmC) [31].

Methylated DNA immunoprecipitation followed by sequencing (MeDIP-Seq) is another widely used method to study DNA methylation profiles. MeDIP enriches methylated DNA fragments by using monoclonal antibodies against 5-methyl cytosine. The immunoprecipitated sample can then be analyzed by high-throughput sequencing to identify the methylated regions and compared to total input DNA that did not undergo immunoprecipitation, thus assessing the enriched fraction representing methylated regions. This technique was also adapted to detect 5-hydroxymethylcytosine (5hmC) DNA [32]. 5hmC, as a different epigenetic modification, is an oxidized form of 5mC [33]. Unlike 5mC, which is associated with transcriptional repression, 5hmC is associated with gene transcription [33]. It is reported that 5hmC may increase gene expression through increasing chromatin accessibility or inhibiting repressor binding [33, 34]. Although MeDIP does not provide information at single-base resolution; it has sufficient resolution to study differentially methylated regions (DMRs), which are functionally more important than single methylation polymorphisms (SMPs) [35-37]. Most CpG dinucleotides are methylated, and there are about 30 million CpGs in the human genome, where 7% of CpGs are located within CpG islands (CGIs). About 70% of annotated genes’ promoters are associated with CGIs, mostly with housekeeping functions [38]. The MeDIP method can target CpG-islands and be enriched for the entire length of the genes [39].

Early MeDIP-Seq methods also required one microgram of input DNA [40]. Several recent protocols use less fragmented DNA per immunoprecipitation (IP) reaction (100 ng to 1 µg) [41-44] but typically require 3-5 days to process [42] for each sample. In those protocols, DNA isolation is performed on individual samples, and DNA is then sheared by sonicators. This process can take *∼*1-2 days (depending on the sample size and desired fragment distributions) [44]. Sonication is followed by immunoprecipitation and library preparation for sequencing. which approximately takes an additional *∼*2-3 days [44]. The ability to process samples in high throughput is limited currently due to the single-sample procedure of the MeDIP step, an issue we address here. We have developed a MeDIP-Seq multiplexing strategy that circumvents these challenges and allows analysis of many samples in parallel, which reduces processing time, cost, and the amount of DNA sample needed. The new platform generated consistent results with as little as 25 ng of DNA that can be isolated from 3400 to 6200 cells [45].

In standard MeDIP, the DNA library construction occurs after immunoprecipitation (IP). In the Mx-MeDIP-Seq approach, the library was prepared before IP, i.e., the IP reaction was sandwiched between index ligation and PCR amplification followed by sequencing. In Mx-MeDIP-Seq, unique barcodes were ligated to DNA from different samples. The prepared DNA libraries were pooled, immunoprecipitated, sequenced, and then demultiplexed. We confirmed the reliability of this Multiplexed MeDIP-Seq (Mx-MeDIP-Seq) protocol by comparing it to standard MeDIP for the same samples individually, including the DNA fold enrichment and the correlation between the two groups after immunoprecipitation. Performing IP reactions with barcode ligated libraries and the ability to combine as many as 10 samples in a single run and optimizing the amount of 5mC antibody could reduce the amount of required starting DNA per sample and reduce time and cost, all prerequisites for developing an automated platform to perform high throughput MeDIP-Seq.

## Methods

### Peripheral Blood Mononuclear Cells (PBMCs) Isolation

To isolate the peripheral blood mononuclear cells (PBMCs), peripheral whole blood was drawn from healthy patients into CPT™ tubes (BD Vacutainer, cat no 362761, Becton Dickenson, Franklin Lakes, NJ). The tubes were centrifuged according to the manufacturer’s directions. The plasma upper layer was removed, and the remaining buffy coat diluted into an equivalent volume of PBS + 2% FBS in 15 mL conical centrifuge tubes and respun. After each centrifugation, the pellet was resuspended into PBS + 2% FBS. Following two washes and centrifugations, the PBMCs were resuspended in Bambanker™ freezing media and stored in liquid nitrogen for future use.

### Micrococcal Nuclease Digestion

The Micrococcal Nuclease Digestion method was based on published methods [46] with some modifications. Frozen PBMCs were centrifuged twice at 600x g for 10 minutes at 20°C and washed with RPMI+ 20% FBS. The cell pellet was resuspended in PBS to achieve a final volume of 7,500 cells per 1µL of suspension. An equal volume of the MNase digestion buffer (0.02 U/µL MNase (Thermo Scientific, Cat. No: 88216) in 2x lysis buffer was added to the cell suspension and gently vortexed. The cell suspension was then incubated on ice for 20 minutes, followed by incubation at 55°C water bath for 10 minutes. The digestion reaction was terminated using 30 mM EGTA. Then the digested chromatin was protease k (*Ambion*™) treated at 55°C and then pre-cleared with 1:2 KAPA Pure beads (Roche Cat. No: 07983298001). The quality and quantity of fragmented DNA were verified using agarose gel (1.5%) electrophoresis and Qubit® 4.0 Fluorometer (Thermo Fisher Scientific, Waltham, Massachusetts, U.S), respectively. The resulting purified DNA was utilized for library construction.

### End-repair, A-tailing, and adapter ligation

Illumina KAPA LTP Library Preparation Kit was employed for library construction (Cat. No: KK8233). Purified, fragmented DNA was end-repaired to produce 5′-phosphorylated, blunt-ended dsDNA fragments. Unique Dual Indexes (UDIs) were ligated to 3’-A-tailed library fragments holding appropriate adapter sequences on both ends. Indexed DNA fragments were subjected to methylated DNA immunoprecipitation, followed by amplification.

### Multiplex Methylated DNA Immunoprecipitation Sequencing (Mx-MeDIP-Seq)

Adapter ligated DNA fractions were pooled together, and methylated DNA immunoprecipitation was performed on the pooled, denatured DNA samples. This method comprises immunocapturing methylated DNA fragments using methylated cytosine-specific antibodies. The affinity of the antibody used in MeDIP enables the detection of any methylated cytosine and is not restricted to the analysis of CpG island. MeDIP was done according to the manufacturer’s instructions (Active Motif Cat. No: 55009). The procedure was optimized based on the input DNA and the quantity of required antibodies. Fragmented DNA was used (50 ng,100 ng, 200 ng, 500 ng, 1 µg) with different amounts of antibody (0.5 to 8 µg). The reaction was carried out in special low DNA LoBind tubes (Eppendorf; Catalog No. 022431048) to minimize DNA loss during the MeDIP procedure. The enriched fraction of the immunoprecipitated (IP) sample DNA was amplified, followed by a real-time quantitative polymerase-chain-reaction (qPCR) technique to analyze the final enrichment for methylated DNA. To confirm the enrichment of methylated DNA in the immunoprecipirtated samples, four primers set specific for the highly methylated region were used as positive controls (*TSH2B, BRDT, ZC3H13, NBR2* genes). *TSH2B* and *BRDT* genes are transcribed exclusively in testis, and CpG sites of this gene are methylated in all somatic tissue. *ZC3H13* is a known positive methylated locus in human cells with no expression in PBMCs [47]. *NBR2* is located within a large CpG island and is found to be methylated in most somatic cells [48]. Similarly, one primer for the unmethylated region (*GAPDH* gene) was selected as the negative control. Normalize Percent Recovery (NPR) was calculated as per Equation 1 and Equation 2, adapted and modified according to this study [42]. Moreover, the fold change in gene expression and fold change ratio were calculated as per Equation 3 and **Error! Reference source not found**..

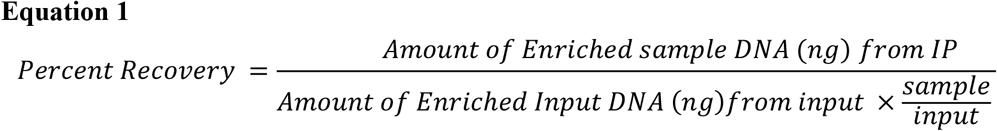

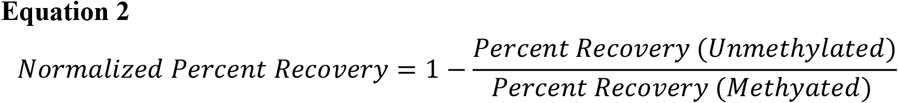

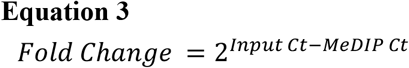

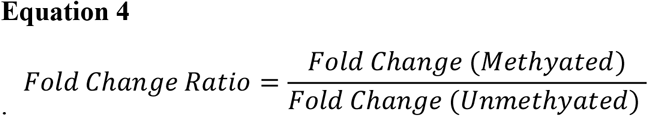

### Size selection

Using KAPA pure beads, library size selection was performed in a two-step process based on manufacturer recommendations. The library fragments ranging between 200 bp and 1000 bp were selected, and the rest were eliminated. First, DNA fragments larger than 1000 bp were removed by incubating the library with 0.5X volume of KAPA pure beads for 5 minutes at room temperature. Second, to capture the fragments above 200 bp, the beads were magnetized, and the supernatant was incubated with 0.5X volume of KAPA beads. After 5 mins of binding at room temperature, the beads were washed 2X with 80% ethanol and resuspended in 10 mM Tris for DNA elution. The supernatant contains the size selected library DNA between 200 to 1000 bp.

### Sequencing

Library quality was analyzed on an Agilent Tapestation and quantified by qPCR (KAPA KK4835-Thermo Fisher Sci Quantstudio 5) at The Arizona State University (ASU) genomic core facility, before pooling and sequencing (2×150 flow cell - Illumina NovaSeq 6000) at the University of Colorado Anschutz Medical Campus Genomics Core Facility.

### Bioinformatics analysis

Cutadapt (v 1.18) [49] was used to remove the adapter sequence, and low-quality ends from raw sequencing reads. Cleaned reads were aligned to the human reference genome (GRCh38.p12, hg38) using Burrows-Wheeler Aligner (BWA v 0.7.17 [50]) to generate BAM files. MethylQA (v 0.2.1) [51] was used to generate genome density for MeDIP-Seq data and reads mapping and CpG coverage statistics. Aligned reads were used to call peaks with MACS2 (v 2.2.6) [52] using input as a control to identify enriched areas in the genome based on performed methylated immunoprecipitation. Next, the bdgcmp utility from the MACS2 package was used to deduct noise by comparing two signal tracks (IP and input) in bedGraph containing the log10 value of fold enrichment. The percentage of CG coverage of cleaned reads was compared to assess the similarity between individual and pooled samples. Next, the correlation test of read counts on common peaks regions for all different samples was performed. Before computing the correlation, all overlapping peaks regions between individual and pooled samples were first merged into one bed file. The bed file contains all regions that should be considered for the coverage analysis. Then multiBamSummary from deeptools2.0 computed the read coverages (the number of unique reads mapped at a given nucleotide) over the merged peaks by using BAM files (individual and pooled samples). Finally, deepTools were used to plot correlation to visualize the multiBamSummary file.

To calculate the enrichment correlation between individual and pooled processed samples on CpG island, we implied ComputeMatrix from deepTools to calculate the log10 value of fold enrichment scores of each base in the CpG islands. In this method, gene body length was divided by the bin size, which was considered 50 in this study. The fold enrichment average in each bin across the CpG islands was then calculated. The common regions overlapping two files were selected per genome region (CpG island here). Pearson correlation was used to calculate the correlation score between the average enrichment scores from the common region in two files.

### Statistical Analyses

Statistical analysis for qPCR data was carried out using one-way ANOVA to compare pooled processed samples and individually processed with different amounts of starting DNA library with respect to Normalize Percent Recovery and fold change ratio, respectively. Tukey method was used for multiple comparisons to determine whether there is a difference between the mean of all possible pairs. Statistical significance is indicated by * for p-values<0.05, ** for p-values<0.01, and **\*\*\*** for p-values**<**0.001.

## Results

### Development of Mx-MeDIP-Seq

The designed workflow to expedite the MeDIP process and minimize the required input DNA is illustrated in Fig. 1**Error! Reference source not found**.A. We elected to fragment the chromatin using micrococcal nuclease to avoid both the time and sample requirements inherent to mechanical fragmentation. Conditions for enzymatic digestion were chosen (**Error! Reference source not found**.) to produce about 30% mono-nucleosomal DNA and an average of 25% di-nucleosomal DNA and minimize DNA loss during library construction. The band percentage was calculated by ImageJ. Bead-based size selection was used to exclude DNA over 1,000 bp and under 200 bp; the remaining digested products were included in the library construction. Removing large fragment sites reduced the noise and increased resolution after sequencing.

**Figure 1.**
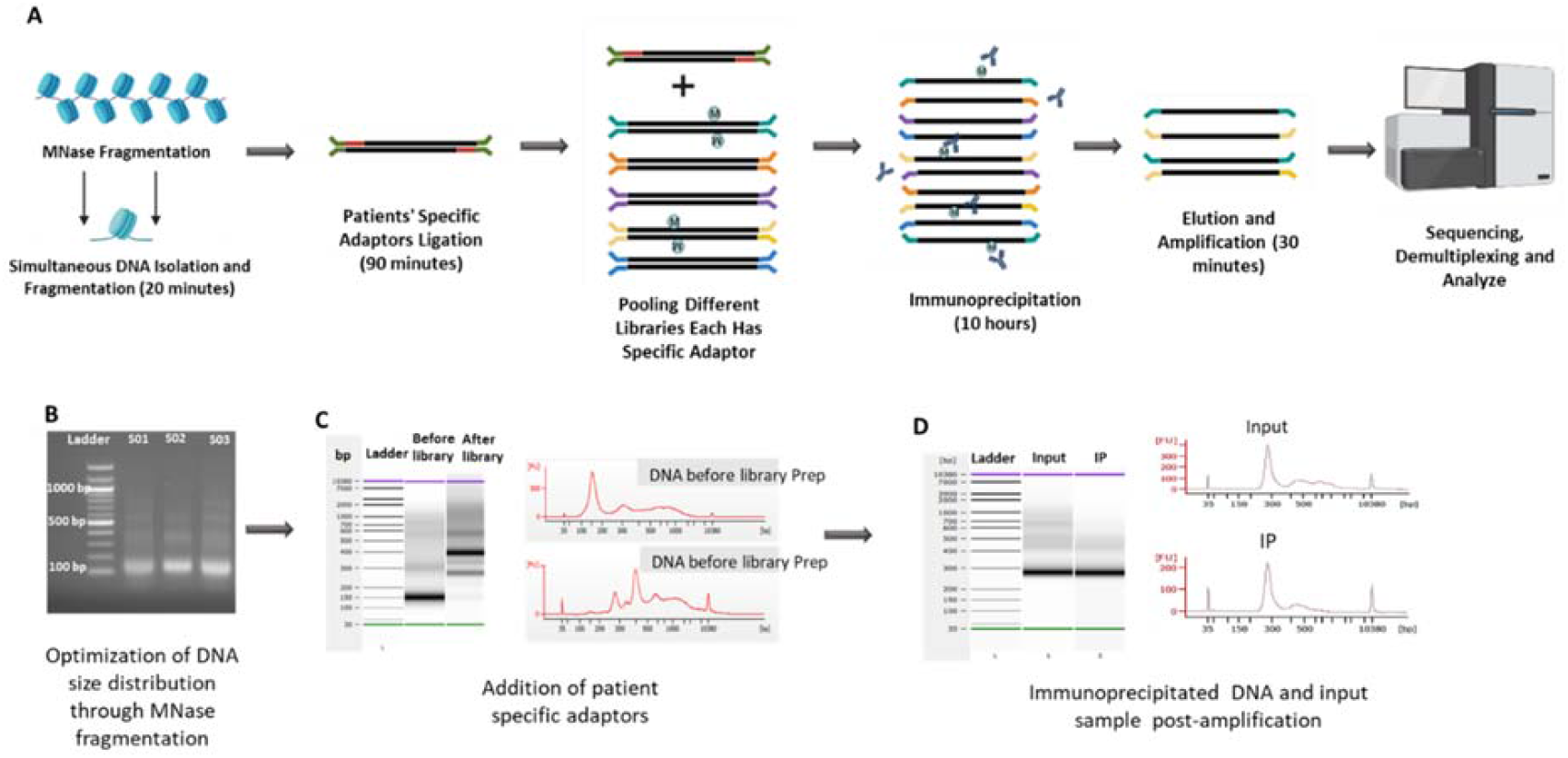
Mx-MeDIP workflow and DNA fragment analysis throughout the experiment. A: Automation friendly high throughput DNA fragmentation, patient specific adapter ligation DNA, immunoprecipitation, and sequencing. B: Representative image of agarose gel (1.5%) electrophoresis for DNA fragmented by micrococcal nuclease (MNase) digestion. The chromatin has been isolated from peripheral blood mononuclear cells (PBMCs). The left lane is a 100 bp marker. MNase digestion conditions yielded about 30% mono-nucleosomes and 25% di-nucleosomes (analyzed by ImageJ). C-Left: Pattern of fragments after library prep and adapter ligation. The fragments shifted by 150 bp due to adding the adapter indexes during library prep. C-Right: Electropherograms of High-Sensitivity chips on an Agilent Bioanalyzer 2100, Pattern of fragments after purification and before library prep (top). The bioanalyzer gel image illustrates the electropherograms result for DNA fragments before (top) and after library prep (bottom). DNA library has proceeded for multiplexing and MeDIP. D-Left: The fragment pattern of DNA after methylated DNA immunoprecipitation and input samples obtained from High-Sensitivity chips on an Agilent Bioanalyzer 2100. E: Electropherograms of High-Sensitivity chips on an Agilent Bioanalyzer 2100, the fragment pattern of DNA after methylated DNA after methylated DNA immunoprecipitation, and input samples

Before pooling different samples, unique patient-specific barcodes were included in the adaptors ligated to the DNA fragments. Unique dual (UD) index adapters from Illumina (Cat# 07141530001) were used, containing 96 different pairing strategies. Adapter ligation was confirmed by the mobility shift of the mono-nucleosomal, and di-nucleosomal bands in the e-gel image and electrogram, as shown in Fig. 1C. A key aspect of this design is that all DNA manipulations steps like end repair, A-tailing and index-ligation are amenable to automation using a standard liquid handling robot.

Multiplexing was achieved by combining several purified DNA libraries after appending unique barcode adapters (up to 10 in this study). In one tube, the mixed library was subjected to methylated DNA immunoprecipitation. Performing this process on all samples simultaneously simplified the workflow by reducing hands-on time, reagents, error, and complexity. It shortened our processing time for 100 samples from one week to one day (Figure 2). The library DNA, magnetic beads, and 5mc-antibody were incubated overnight. The amount of antibody used was based on the total amount of library DNA (Fig. S2). The enriched DNA was then PCR amplified and purified, removing DNA over 1,000 bp by size-selection with Ampure XP beads. Larger fragments (> 1000bp) were shown to have lower efficiency and yield during amplification on the flow cell surface while sequencing [53, 54]. After IP, we confirmed that the resulting DNA contained only fragments between 300 bp to 1000 bp on an Agilent 1200 bioanalyzer (Figure 1B).

**Figure 2.**
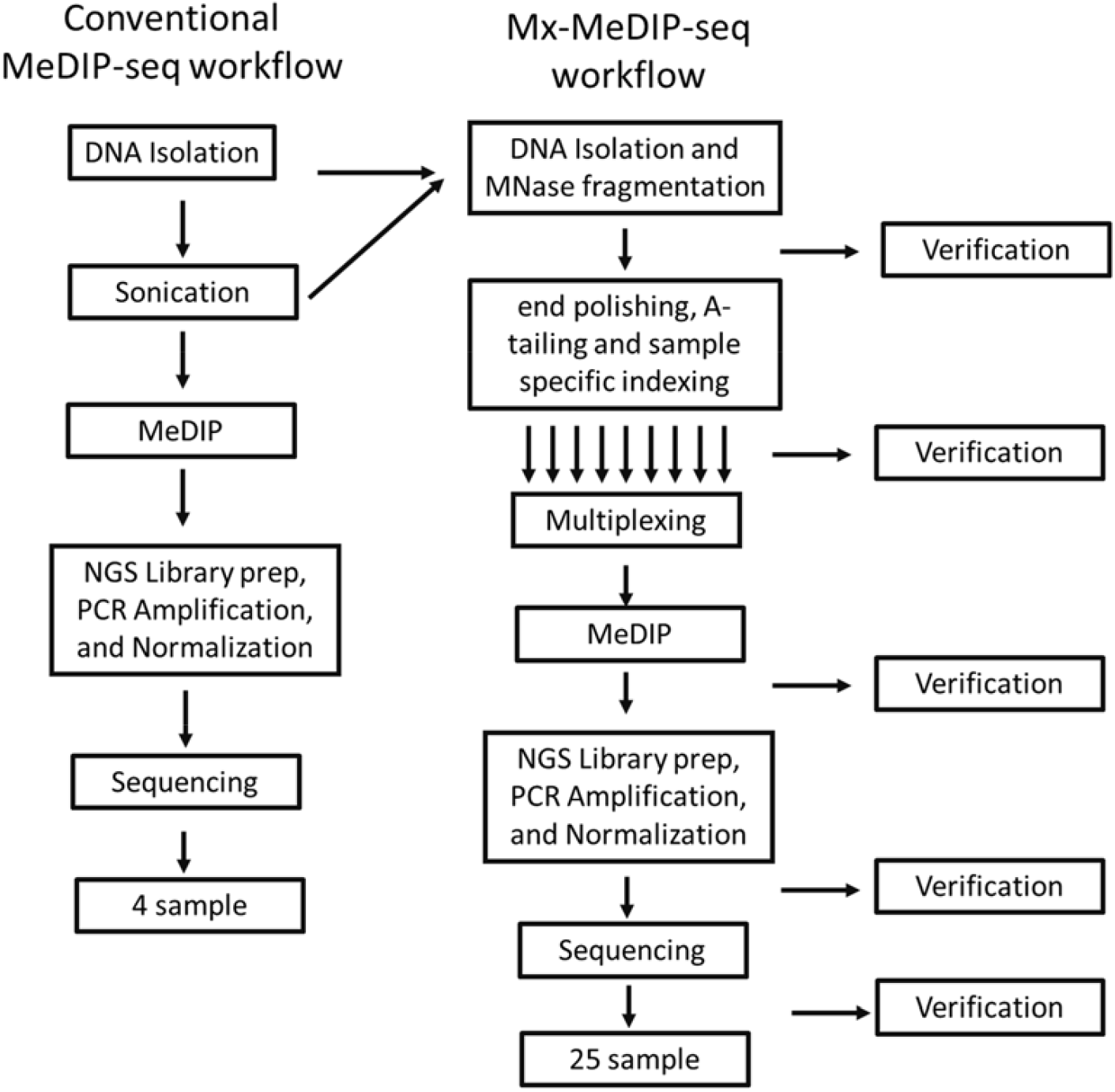
Mx-MeDIP-Seq workflow in comparison to standard workflow. Mx-MeDIP-Seq combines two steps for DNA isolation and fragmentation into one step. Immunoprecipitation is sandwiched between two steps of library preparation. In the first step, each sample gets a unique adapter. Several samples can be put together, followed by immunoprecipitation. The isolated methylated DNA is amplified and sent to sequencing, followed by demultiplexing and analysis.

Several primer pairs were selected for enrichment estimation based on qPCR, which amplifies either methylated or unmethylated DNA fragments (see Table 2, primer pairs 2 for sequences). *TSH2B, BRDT, ZC3H13*, and *NBR2* genes were selected as the positive controls since they are highly methylated in PBMC cells [47, 48, 55-57]. The human *GAPDH* primer was considered a negative control that recognizes the human *GAPDH* promoter as an unmethylated region in all somatic cells [55, 56, 58]. The efficiency of the immunoprecipitation in the MeDIP was assessed based on Ct qPCR data using normalized percent recovery and fold change ratio formula adapted from the manufacturer’s protocol and published studies [42, 55, 56] (Equation 1 to **Error! Reference source not found**.). A MeDIP reaction was passed on for DNA sequencing when there was a high recovery (normalized percent recovery ≥ 95%) of known methylated over unmethylated fragments and a fold change ratio >5 [59, 60].

**Table 1:** 2x Lysis buffer stock preparation contacting MNase. This mixture is used to lyse the cells and isolate and fragment the DNA strand in one step.

| Reagents | Volume (mL) |
| --- | --- |
| 100mM Tris-HCl (1 M, pH 8.0) | 20 |
| 300mM NaCl (5 M) | 12 |
| 2% Triton X-100 (25%) | 16 |
| 0.2% sodium deoxycholate (DOC, 12.5%) | 3.2 |
| 10mM CaCl<sub>2</sub> (1 M) | 2 |
| 10 mM sodium butyrate (800mM) | 2.5 |
| H<sub>2</sub>O | 144.3 |
| 1 U/μL MNase Stock solution | 4 |
| **Total Volume** | **200** |

**Table 2.**
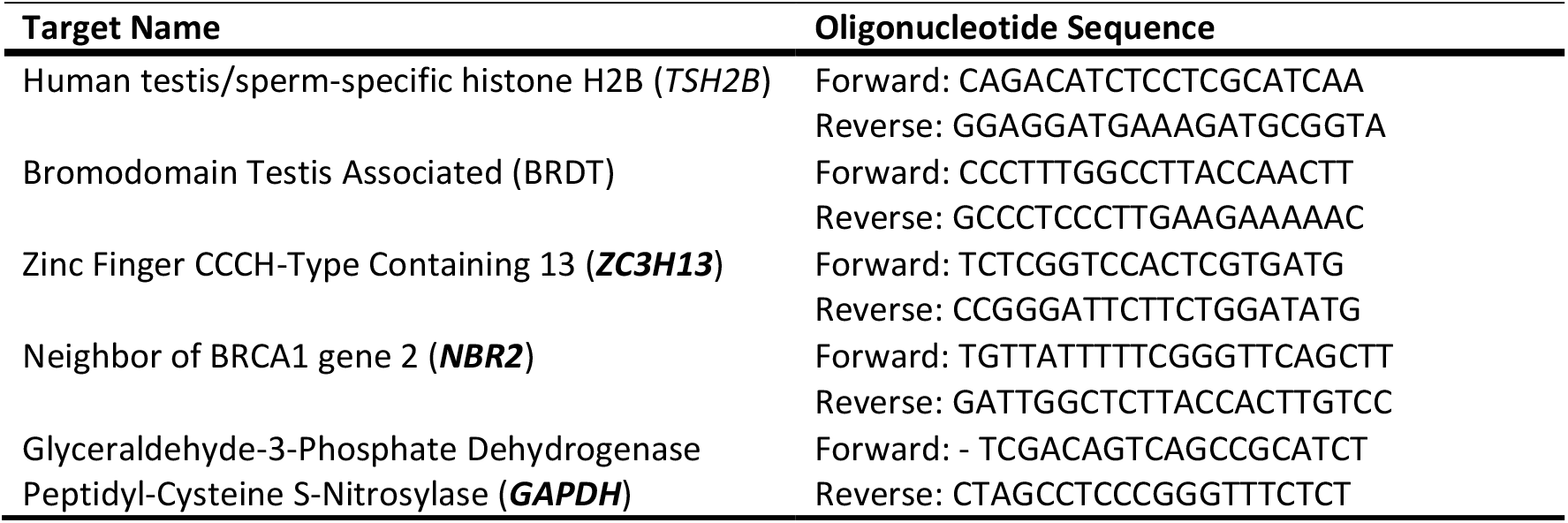
Primers used in qPCR to assess the enrichment of pooled down DNA. The first four primers were considered positive controls, and *GAPDH* was considered a negative control

Sample comparisons were made by first establishing sample quality by examining %CG coverage after demultiplexing and adaptor sequence removal (Figure 3). Then reads on common peaks and enrichment of CpG islands were correlated and compared.

**Figure 3.**
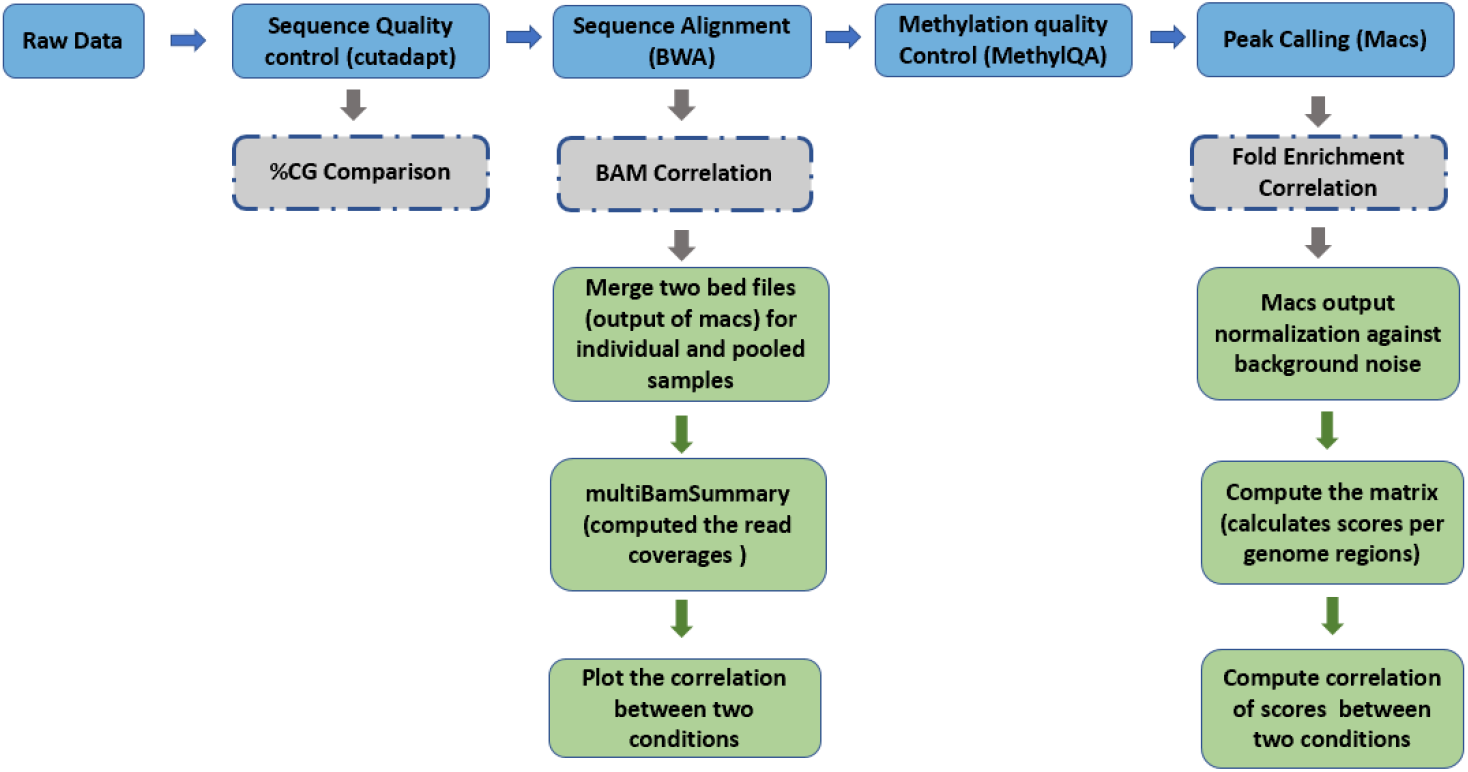
Stages and component steps of bioinformatics data analysis. Sequence data from MX-MeDIP-Seq and MeDIP-Seq were compared in three steps by comparing %CG, genome-wide read coverage correlation, and fold enrichment correlation after peak calling.

### Sonication vs. enzyme fragmentation on methylation data

In order to compare fragmentation by MNase and sonication on methylation result, three MeDIP-seq data were downloaded from Gene Expression Omnibus under accession number GSE39604 [61]. In this study, PBMCs were isolated from freshly collected blood and sonicated to approximately 100–500 bp. Methylated DNAs were enriched by immunoprecipitation with an anti-5-methyl-cytosine monoclonal antibody similar to our study. The intersect of each MedIP-seq data set on CpGIsland was calculated using bedtools/2.30.0 intersect command. Out of 30,344 CpGIsland listed in the human genome, the enzymatically fragmented MeDIP-seq matched 26234 ± 3765 sites, while groups with sonicated fragments matched 24941 ± 4591 sites. These two samples matched on 89.8% ± 7.6% of sites on CpG islands.

### Individual MeDIP and Mx-MeDIP Resulted in Similar Enrichment and Correlation

To investigate whether multiplexing affected the enrichment and sequencing results, a set of ten different library DNAs with unique adapters were tested individually or as a single pool using standard MeDIP-Seq [41, 44, 62] (Figure 4A). The enrichment for every sample was assessed with qPCR vs. input samples (Figure 4B and C).

**Figure 4.**
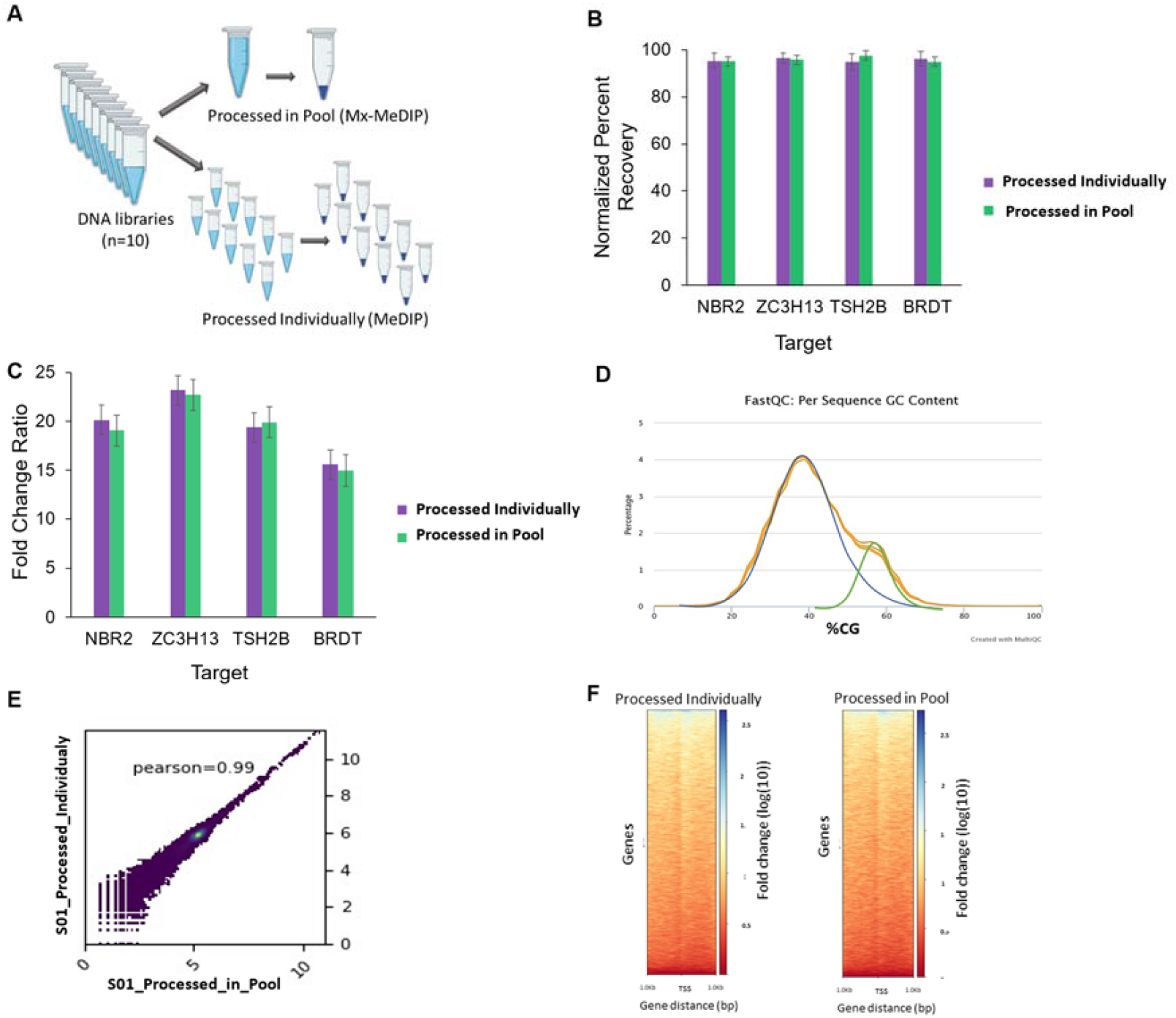
Comparison of samples processed in a pool and same samples processed individually before and after sequencing. A: illustration of the difference between processed individually group (MeDIP) and processed in pool group (Mx-MeDIP). B and C, qPCR result: Normalized percent recovery (B) and fold change ratio (C) for samples that were processed in the pool (Mx-MeDIP-Seq) and the same samples processed individually (standard MeDIP-Seq). Ten different library DNA went through MeDIP individually. Also, a pool containing all those ten samples went through IP simultaneously. The plots show a qPCR result analysis for individual (n=10) and pooled samples. D: empirical GC content in the MeDIP enriched samples (orange line) compared to the theoretical distribution (blue and green line) E: representative images of correlation of read coverage between individual and pooled samples (rest of correlation plots illustrated in Figure S1. F: Representative heatmap plot of peaks called on CpG islands for a sample processed individually and samples processed in the pool (detailed information in Table 3)

As shown in Figure 4B, the average normalized percent recovery was over 95% for all the assessed targets. Samples that were processed individually showed similar normalized percent recovery compared to samples in the pool. There was no significant difference between individual and pool processing, indicating that multiplexing did not affect the measured normalized percent recovery. The fold change ratio vs. input was also similar for each target (*NBR2, ZC3H13, TSH2B, BRDT*) for samples in both groups. The plot shows that the fold change ratio was between 15 ± 1.6 for *BRDT* and 24 ± 1.8 for *ZC3H13* (mean ± SD; Figure 4C). The sequencing of the methylated enriched DNA yielded approximately 43 ± 9.9 (mean ± SD) million reads per sample with the mean Phred scores above 35 in both pooled and individual assays. The percent of GC content on obtained sequencing reads was measured using FastQC. Since the samples were immunoprecipitated based on GC-rich DNA (i.e., CpG islands), we expected to see a shifted distribution to the right side of the theoretical genomic GC distribution. Figure 4D shows that there was a clear enrichment of GC-rich reads in the MeDIP samples, and the shifted distribution was similar for both individual and pooled groups. The quality control analysis also revealed that both groups (individual and pooled) had similar % GC content (40.3% ± 0.24% vs. 40.2% ± 0.8% (mean ± SD)) and shifted the distribution of %GC (59.1% ± 0.70% vs. 59.1% ± 0.53% (mean ± SD); Table 3).

**Table 3.** Summary of GC content and correlation coefficients between ten samples that were processed individually (conventional MeDIP-Seq) and the same sample processed in the pool (Mx-MeDIP-Seq)

| **Samples** | **Shifted peak in %CG (Processed individually)** | **Shifted peak in %CG (processed in pool)** | **Pearson correlation coefficient (of BAM files between two groups)** | **Pearson correlation coefficient (of Peaks called on CpG island)** |
| --- | --- | --- | --- | --- |
| S1 | 59 | 59 | 0.99 | 0.998 |
| S2 | 60 | 59 | 0.99 | 0.996 |
| S3 | 59 | 59 | 0.90 | 0.996 |
| S4 | 59 | 60 | 0.89 | 0.994 |
| S5 | 59 | 60 | 0.89 | 0.998 |
| S6 | 58 | 59 | 0.88 | 0.996 |
| S7 | 60 | 59 | 0.87 | 0.999 |
| S8 | 59 | 58 | 0.87 | 0.995 |
| S9 | 58 | 59 | 0.89 | 0.993 |
| S10 | 60 | 59 | 0.99 | 0.993 |
| Mean | 59.10 | 59.10 | 0.91 | 0.995 |
| standard deviation (SD) | 0.70 | 0.53 | 0.05 | 0.002 |

The overall similarity of read coverages between the pooled and the individually processed samples was then assessed. When the Pearson correlation coefficients (R-squared) of the read counts (*i*.*e*., the number of unique reads mapped at a given nucleotide) across the genome and within the called peaks were calculated, the read coverage between these two groups was highly correlated, as shown in a representative image in Figure 4E, and Fig. S2 (supporting information) illustrates the correlation plots for other samples. Overall, the average correlation coefficients between the read coverages for individual and pool processing were 0.91 ± 0.05 (mean ± SD; Table 3), demonstrating a high similarity of genome-wide read coverage correlation and read coverage on merge of all samples’ peak regions for both samples of both groups.

Fold enrichment per CpG island was also evaluated between individual and pool processing methods. The Pearson correlation coefficient between the fold enrichment values for every bin in the gene body for the two groups (individual vs. pool possessing, Figure 4F) was calculated and summarized in Table 3.

. For all the samples, the result of ComputeMatrix over CpG island showed a strong correlation (Pearson correlation coefficient always > 0.95), indicating each sample either processed through Mx-MeDIP or standard MeDIP have peaks called on a similar region or CpG island.

### Optimizing Sample Needs for Mx-MeDIP-Seq

As it is not always possible to get identical yields from each sample, a potential concern is whether a greater or lesser representation of individual samples in the pool could affect the outcome. To test this notion, several pools were prepared containing 200 ng of input DNA but containing different ratios of constituent samples. One pool contained an equal distribution (25%) of each sample (50 ng each, Figure 5A). The second pool contained different amounts of each DNA sample; the first sample was 10% of the pool (20 ng), the second sample was 20% (40ng), and samples 3 and 4 were 30% (60 ng) and 40% (80 ng), respectively (Figure 5A). After three Mx-MeDIP repeats, normalized percent recovery and fold enrichment were assessed using qPCR, and read depth and fold enrichment per CpG islands were assessed using sequencing. Enrichment efficacy evaluation through qPCR (Figure 5B) showed that normalized percent recovery was > 95% for all four positive targets. These two pools had similar fold enrichment ratios of 21.6 ± 2.1 vs. 21.2 ± 2.2 (mean ± SD, equally-distributed and differentially-distributed pools, respectively). The results suggest that since each pool contained the same amount of DNA (200 ng), the final enrichment and normalized percent recovery were similar, independent of the amount and proportion of each DNA sample.

**Figure 5.**
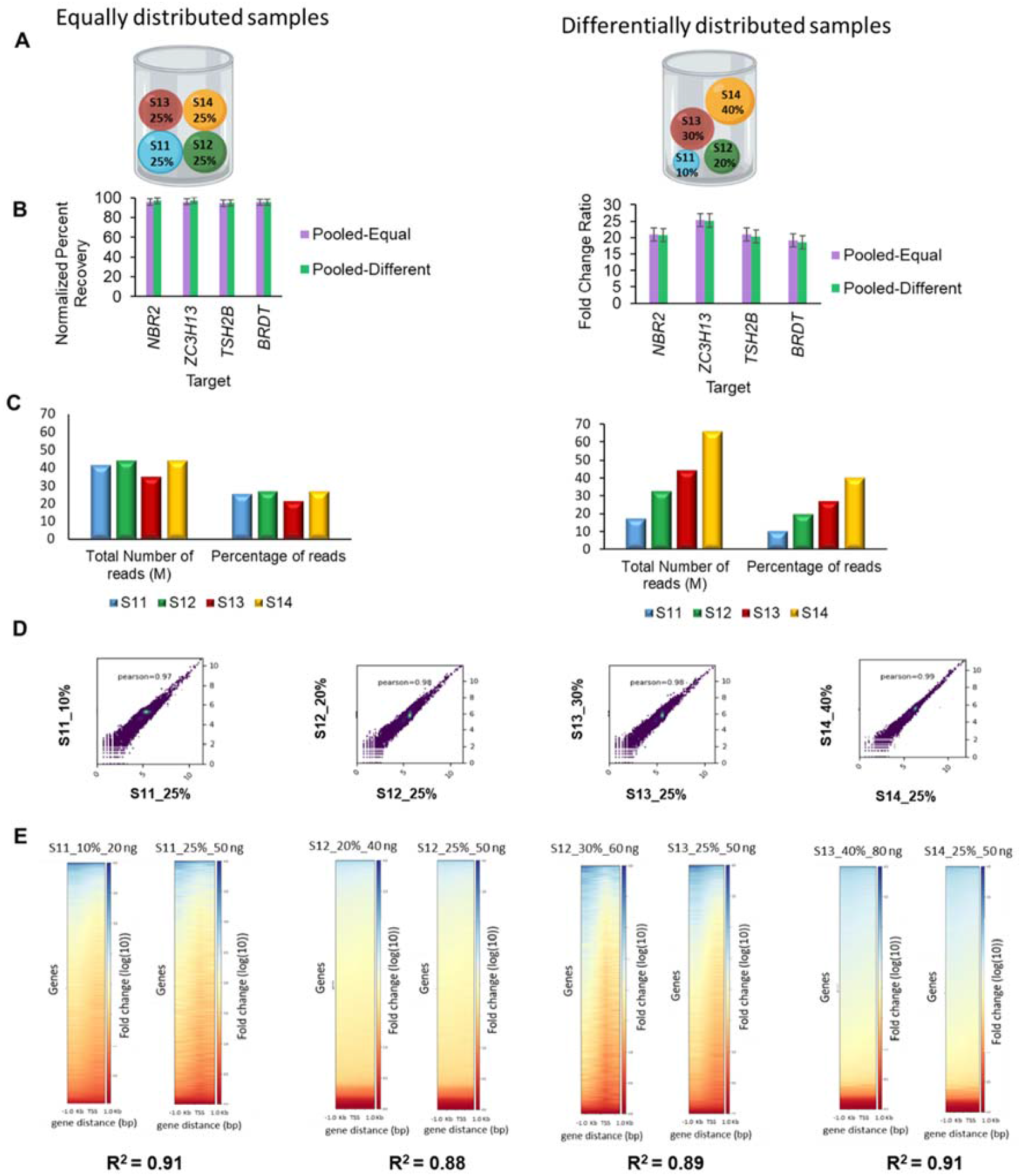
Comparison of Mx-MeDIP-Seq pools with equal distribution and different distribution of DNA. A: pools that contain equal amounts (25% each, equally distributed samples) vs different amounts of DNA samples (ranging from 10% to 40%). B: qPCR result Normalized percent recovery (left) and fold change ratio (right) for a pool of equal amounts of starting DNA vs. the different amounts of starting DNA. For these studies, we carried out n=3 for each condition. C: total number of reads (million) and percentage of reads of each sample in the equally distributed and differentially distributed pool after sequencing. D correlation of reading coverage between equally and differentially distributed pool. Left to right (sample_11 (S11) with 10% in a differentially distributed pool and 25% in an equally distributed pool. S12 with 20% in the differentially distributed pool and 25% in the equally distributed pool. S13 with 30% in the differential pool and 25% in the equally distributed pool. S14 with 40% in the differential pool and 25% in the equally distributed pool. E: Comparing the peak correlation among samples with different amounts of initial input DNA for Mx-MeDIP-Seq (left to right: Heatmap plot of peaks called on CpG islands for S11 with 20 ng initial DNA compared to 50 ng of initial DNA. S12 with 40 ng initial DNA compared to 50 ng of initial DNA. S13 with 60 ng initial DNA compared to 50 ng of initial DNA. K. S14 with 80 ng initial DNA compared to 50 ng of initial DNA).

Similarly, the percentage of sequencing reads for each sample corresponded directly to the fraction that sample comprised in the total pool (Figure 5C). For Pool_1, comprising 4 samples equally distributed, the percentage of total reads after sequencing ranged from 21% to 27% for each sample. On the other hand, pool_2 comprised a different fraction of each sample; the sequencing read percentage (11.6%, 21.9%, 27.7%, and 38.8%) corresponded closely to the percentage of each sample in the pool (10%, 20%, 30%, and 40%) (Figure 5C).

To assess the similarity between read coverages of samples in the equally distributed pool and the differentially distributed pool, scatter plots were acquired using the deeptools package (Figure 5D). The read coverage correlations between equal and differentially distributed pools were 97% to 99% for all four samples, indicating that the qPCR and sequencing results were insensitive to differences in the fraction that a sample comprises in the pool. Next, the peaks located on CpG islands were evaluated. The Pearson correlation coefficient between sample making 10% of the pool (S11_10%) and sample making 25% of the pool (S11_25%) as well as 20%, 30%, 40% vs. 25%, were calculated. There was more than 88% agreement between the fold change of the peaks called on CpG island while the samples range from 10% to 40% compared to constant with as little as 20 ng of DNA (Figure 5E). As the availability of DNA might vary in different situations, Mx-MeDIP outcomes such as read coverage and fold changes of peaks remained resilient to different relative fractions of DNA material in the pool over a 4-fold range (10 – 40%).

### Mx-MeDIP-Seq Can be Performed Using Small Amounts of DNA

We also investigated how much DNA was required for consistent results. The protocol was tested on three replicates of different amounts of pooled DNA pool, 4 ng, 20 ng, 40 ng, 100 ng, 200 ng, and 400 ng. Each pool contained four different DNA samples with an equal amount of uniquely indexed DNA libraries. After immunoprecipitation, the products were analyzed quantitatively using qPCR to determine the normalized percent recovery and fold enrichment ratios. Mx-MeDIP-Seq was considered successful when normalized percent recovery was ≥95%, and the fold change ratio was >5.

The yield of captured DNA increased as expected with the increased DNA starting material. As Figure 6 indicates, performing Mx-MeDIP-Seq using 25 ng, 50 ng, and 100 ng led to normalized percent recovery ≥95% and fold change ratio >5 for all targets. Moreover, there was no significant difference in normalized percent recovery using 50 ng and 100 ng of starting DNA. Lowering the DNA amount to 1 ng reflected the limitation in distinguishing methylated and unmethylated DNA.

**Figure 6.**
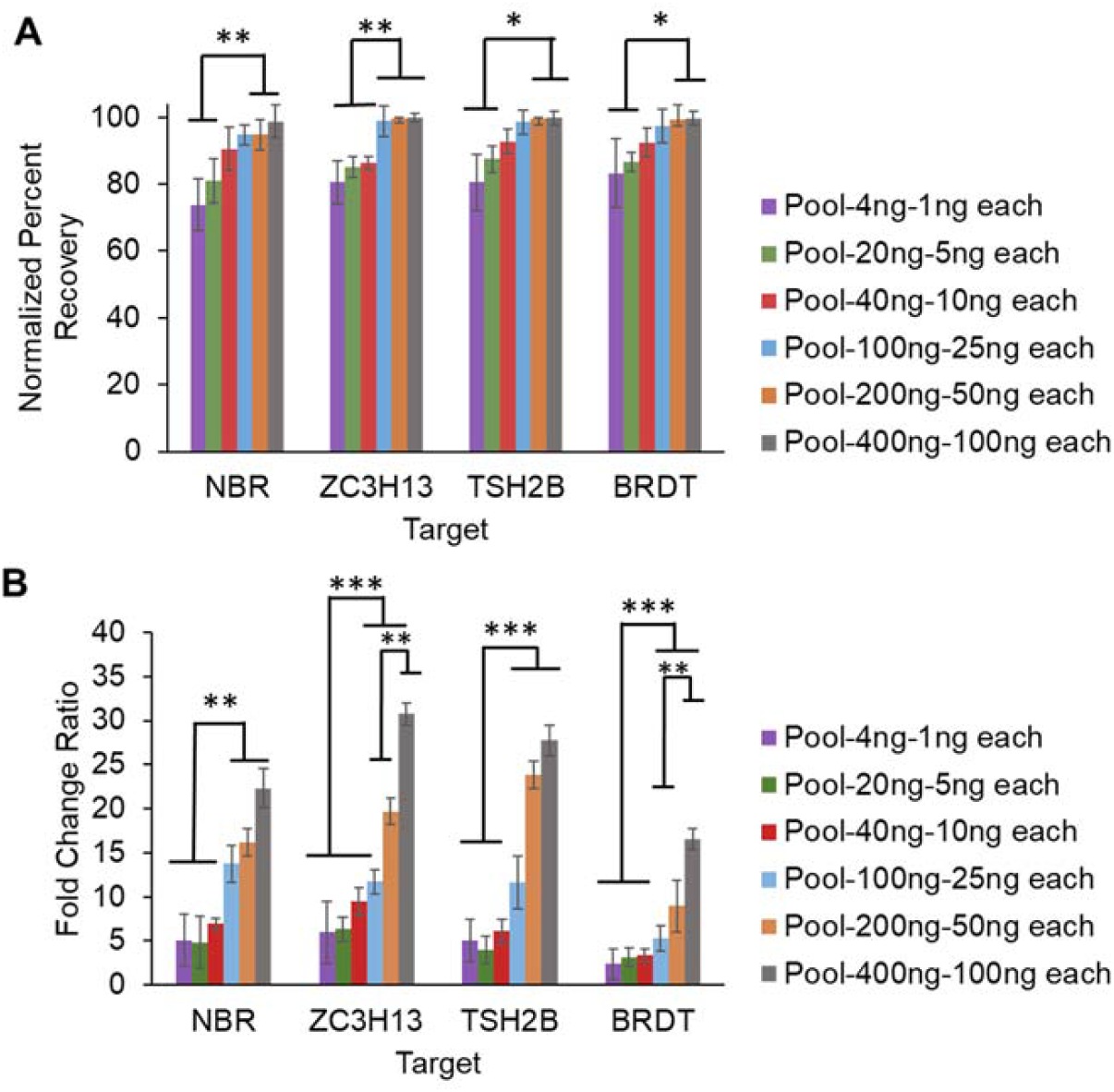
Effect of minimizing the amount of input DNA for Mx-MeDIP-Seq. Normalized percent recovery and fold change ratio for a pool of DNA carried through Mx-MeDIP. A Triplicate of six pools of DNA was prepared. Each pool contains four different DNA samples (the final amount of DNA were 4 ng, 20 ng, 40 ng, 100 ng, 200 ng, 400 ng). Starting DNA amounts were reduced from 100 ng to 1 ng in each different pool. The final enrichment efficacy was assessed using qPCR data. Normalized percent recovery and Fold change ratio hold their high value while the DNA amount was reduced to 25 ng. A significant decrease in Normalized percent recovery and fold change ratio was observed when the DNA amount was reduced to 10 ng. Statistical analysis between groups was determined using one-way ANOVA and Tukey test; p-values < 0.05 were considered statistically significant. For these studies, we carried out n=3 for each condition, and in the figure, *: p-value<0.05, **: p-value<0.01, ***: p-value<0.001.

In another similar experiment (Fig. S2), the Mx-MeDIP technique with different amounts of DNA pools was compared to standard MeDIP using 50 ng of individual DNA samples. The pools contained 50 ng, 250 ng, and 500 ng of DNA composed of ten samples with equal distribution. There was no significant difference using 50 ng of DNA processed individually or in the pool based on enrichment efficiency measurement using fold change ratio and normalized percent recovery. The reduction of starting materials from 50 ng to 25 ng did not show a statistically significant change in normalized percent recovery and fold change ratio. The normalized percent recovery was ≥ 95%, and the fold change ratio was >10 for both groups indicating that lowering the DNA amounts to 25 ng did not compromise the enrichment result. (Figure 6). These two experiments indicated that using 25 ng and 50 ng of individual DNA samples in the pool resulted in successful Mx-MeDIP regardless of the number of samples in the pool in our experimental setting. Additionally, there was a high read coverage correlation and peak correlation of sequence data acquired from 25 ng and 50 ng of DNA (Figure 5D and E).

## Discussion

Epigenetic phenomena are heritable changes in cellular phenotype that are not due to mutations in DNA sequence. DNA methylation is a key epigenetic mark that is associated with transcriptional repression during development, maintenance of homeostasis, and disease [63-65]. Many technologies have been successfully used over the years to measure DNA methylation, each with its own advantages and disadvantages[66]. Bisulfite sequencing method is a treatment that distinguishes unmethylated from methylated cytosines. Common affinity enrichment strategies include methyl-DNA immunoprecipitation using methylation specific antibodies (MeDIP[54, 67, 68]) and methyl-CpG binding domain protein (MBD [69, 70]). Both MeDIP- and MBD-based methods have the advantage of being low-cost and straightforward for laboratories that are already skilled in chromatin immunoprecipitation sequencing (ChIP-seq) or related techniques. The disadvantages include relatively low resolution, susceptibility to copy number variation bias, and a bias toward observations of methylated DNA compared with bisulfite-based protocols[66]. One caveat with affinity approaches is that methylated CpG-rich sequences may give higher enrichments than methylated CpG-low sequences. Improving the capacity and efficiency of the methods available for studying the methylome can expand the study of epigenetics.

Although methylated DNA immunoprecipitation followed by sequencing (MeDIP-Seq) was introduced to reduce the cost and complexity of the data associated with whole-genome bisulfite sequencing methods as the primary method to analyze global methylation information [71], its cost and low throughput remain a barrier to researchers wishing to carry out large-scale genome-wide methylation studies. There are three main barriers for converting low throughput MeDIP-Seq into a high throughput method; 1) DNA purification; 2) un-biased DNA fragmentation; and 3) labor-intensive process of individual immunoprecipitations. In this study, we addressed all three impediments by coupling DNA fragmentation with cell lysis/purification and barcoding individual samples and multiplexing by pooling samples, Mx-MeDIP-Seq allowed individual researcher to process up to sixty samples simultaneously while they could process 6 samples using historical methods.

In Mx_MeDIP-Seq, the use of enzymatic DNA digestion, instead of mechanical fragmentation, provided a uniform size distribution for many samples at once and minimized sample loss during library preparation. DNA over 1,000 bp was removed by size selection with Ampure XP beads. Larger fragments (> 1000bp) were shown to have lower efficiency and lower yield during amplification on the flowcell surface while sequencing [53, 54]. MNase digestion based MeDIP-Seq method produce similar results in identifying CpG islands compared to DNA fragmentation by sonication.

Following purification, the efficiency of the MeDIP reaction was assessed using quantitative PCR of known hypermethylated regions. To define a reasonable criteria threshold, we have found normalized percent recovery to be more than 90% while the fold change ratio ranged from 4.6 to 100 in independent studies [42, 59, 60]. Samples that passed this QC threshold, were taken forward for sequencing. Primer pairs were selected for enrichment estimation based on qPCR, amplified either methylated or unmethylated DNA fragments. *TSH2B, BRDT, ZC3H13, NBR2* genes were selected as the positive controls since they were known to be highly methylated in PBMCs cells [47, 48, 55-57]. The human *GAPDH* primer was considered a negative control that recognizes the human *GAPDH* promoter as an unmethylated region in all somatic cells [55, 56, 58]. Our results show that both approaches (standard MeDIP-Seq and Mx-MeDIP-Seq) resulted in near-identical enrichment efficiency using fold-change ratio and normalized percent recovery criteria (Figure 4B-C), based on known hypermethylated region. It is possible that not every methylated region could yield similar enrichment between conventional and Mx-MeDIP-Seq methods, or between Mx-MeDIP-Seq and bisulfite sequencing.

Poor or inefficient adapter ligation, low starting DNA quality, and quantity, the short incubation time for IP reactions, PCR failure, and inaccurate calculation of normalized concentrations of samples may result in low normalized percent recovery and a low fold change ratio for both MeDIP-Seq or Mx-MeDIP-Seq. However, in the current study, optimizing each step, the normalized percent recovery and fold change ratio hold at success level. Additionally, the manual library preparation step in Mx-MeDIP-Seq can be automated to reduce errors during pipetting, processing time and labor-intensiveness, and produce consistent results for multiple samples.

After purification and amplification, the samples were sent to a next-generation sequencing platform to investigate the DNA methylome. The sequence reads were then demultiplexed based on their unique adapters. This positively affects cost, such that MeDIP-seq becomes at least 10X more cost-effective.

Looking at the sequence results, the read coverage strong correlation (Pearson value of 0.9; Figure 4) was achieved for the multiplexing approach compared to standard MeDIP. A similar correlation analysis was done across the CpG islands, to look at the genome-wide concordance between Mx-MeDIP and standard one, which resulted in a high correlation on a genome wide level, (0.99 for Pearson correlation coefficient) for all ten samples taken through the two approaches. This strong correlation indicates that Mx-MeDIP can produce the same data from large numbers of samples simultaneously and relatively easier. This approach cuts hands-on time by approximately two-thirds compared with conventional MeDIP.

The available yield of individual DNA for Mx-MeDIP-Seq may vary in different situations due to sample availability or due to differences in DNA extraction efficiency. We tested the tolerance of Mx-MeDIP for varying amounts of individual DNA by pooling DNA at different amounts. Mx-MeDIP outcomes such as enrichment efficacy and sequence data showed resilience to using different fractions of DNA material in the pool (Figure 5, Figure 6). A significant correlation (R^2^ > 0.9) for CpG islands between using 20 ng of DNA in a 200-ng pool compared to using 50 ng in a 200-ng pool indicates that Mx-MeDIP produced similar data considering this change. In the case of limited availability of clinical samples, a flexible amount of DNA as small as 20 ng will still give a similar methylome result and excellent normalized percent recovery and fold change ratio. This amount of DNA can be isolated from as few as 3400 to 6200 cells, depending on the tissue type [45], making the method more applicable to precious samples. In terms of throughput, we currently and routinely perform Mx-MeDIP for 100 samples in 24 h with one technician, whereas standard methods [41, 44, 62] can take weeks and cost substantially more.

Although this method can be automated to perform hundreds of samples simultaneously using a simple liquid handling robots such as Beckman i7, we have not shown any results using such methods. We are optimizing methods to perform large number of samples using Beckman i7 robots equipped with different temperature incubation stations, magnetic bead retrieving capabilities and shakers. Experiments covering broad range of DNA concentrations, 25 ng to 2 µg will also be necessary to mitigate varying DNA extraction efficiency of hundreds of patient samples, in a typical clinical specimen pool. We recommend optimizing DNA extraction efficiency for each specimen types.

Multiple automated methylated DNA immunoprecipitation sequencing protocols, such as AutoMeDIP-Seq, GBS-MeDIP, were proposed and proven to work for low to medium throughput sample numbers[60, 72]. The advantage of MeDIP is that genomic capture is relatively unbiased and not limited to restriction sites or CpG islands. GBS-MeDIP uses similar multiplexing strategy to Mx-MeDIP-Seq using *pst1* restriction digestion and adapter ligation[72-74]. GBS-MeDIP have been adapted to multiple tissue/specimen types[73-75]. MxMeDIP provides significantly higher throughput capabilities due to simultaneous DNA fragmentation and purification through MNase digestion. At a resolution determined by DNA fragmentation (110 bp, here), MeDIP-seq offers comparable coverage at much lower cost than MethylC-seq.

In summary, Mx-MeDIP-seq would be the method of choice when a cost-effective, unbiased method requires a small amount of DNA, and which will allow processing of large numbers of samples in one experiment is needed. This method would be the ideal starting point for an automated or even portable device for monitoring epigenomic methylation.

## Conclusion

This study proposed a novel technique to perform methylated DNA immunoprecipitation followed by sequencing (Mx-MeDIP-Seq). DNA was simultaneously extracted and fragmented through MNase digestion. After adapters ligation, different samples with unique adapters were pooled together and underwent immunoprecipitation using an anti-5-methylcytosine monoclonal antibody. We performed Mx-MeDIP-Seq for pools containing 4 to 10 different samples in our study. The result showed that pool processing had similar enrichment efficacy compared to individual processing. Additionally, there was a strong correlation between read coverage and a strong correlation between fold change of peaks on CpG islands of individual and pooled processed samples. Moreover, using a different fraction of DNA in a pool would not affect Mx-MeDIP outcomes such as read coverage and fold changes of peaks. The method was optimized to use different amounts of starting materials for each sample and pool, and 25 ng of DNA showed comparable results with 100 ng of DNA. Altogether, the proposed approach for methylated DNA immunoprecipitation (Mx-MeDIP-Seq) results in a faster and more cost-effective method. It requires fewer amounts of starting DNA and is less prone to technical errors, making this method more applicable to a broader range of samples and studies.

## ACKNOWLEDGMENTS

We sincerely thank Ms. Thiruppavai Chandrasekaran, Ms. Candyce McDaniel, and Ms. Rowan Johnson, research associates, for their support. This research was supported by DARPA (HR001118S0023, AWD00033689) and ABOR (2950007-01).

