## Supporting tables and figures for "Multiplexed Methylated DNA Immunoprecipitation Sequencing (Mx-MeDIP-Seq) to Study DNA Methylation Using Low Amounts of DNA"

Table S1. Preparation of stock reagents used in lysis buffer

| **Stock Reagents** | **Molecular Weight (g/mol)** | **Weight (grams)** | **the volume of water (mL)** |
| --- | --- | --- | --- |
| 5M NaCl | 58.44 | 11.69 g | 40 |
| 1 M Tris-HCl pH 8.0* | 121.2 | 4.8 g | 40 |
| 0.5M EDTA | 292.24 | 5.85 g | 40 |
| 12.5% sodium deoxycholate | 414.56 | 5 g | 40 |
| 10% SDS | NA | 4 mL | 36 |
| 1 M LiCl (Store at 4°C) | 42.394 | 1.7 g | 40 |

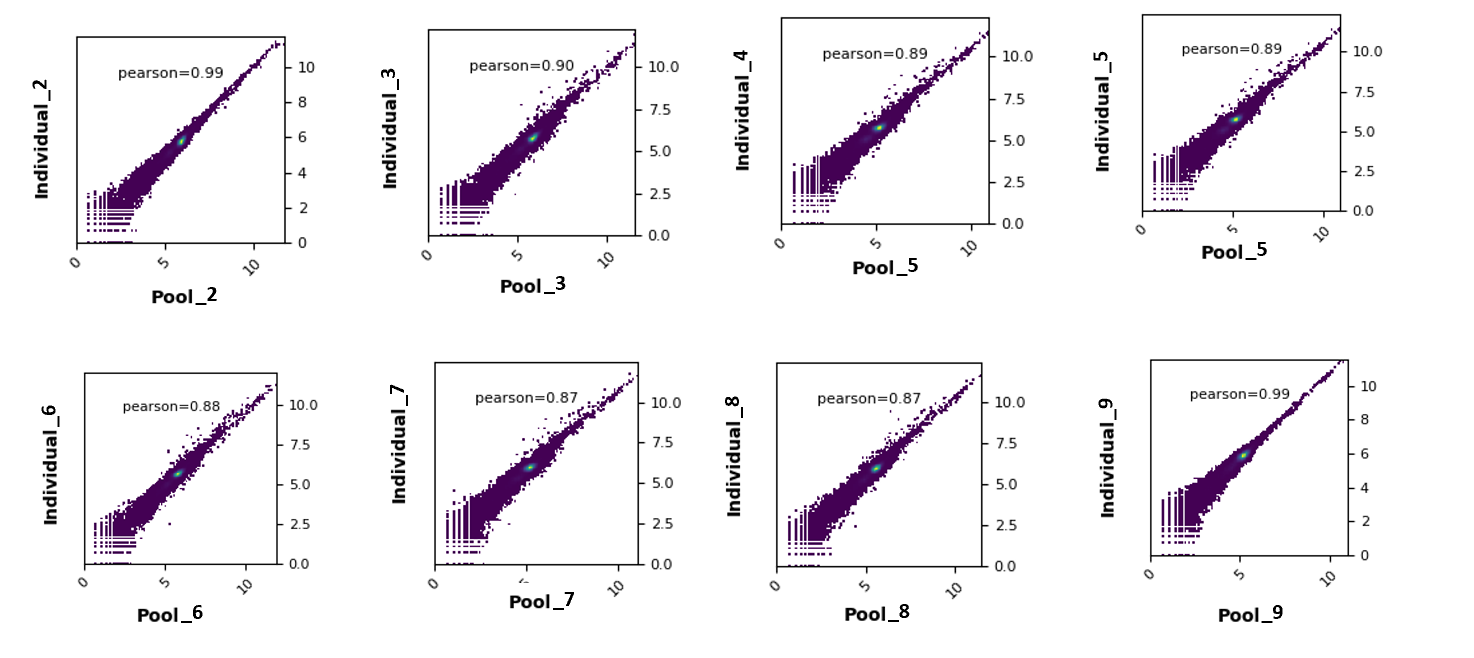

Figure S1. Correlation of read coverage between the samples that were processed in the pool (Mx-MeDIP-seq) and the same sample that was processed individually (conventional MeDIP)

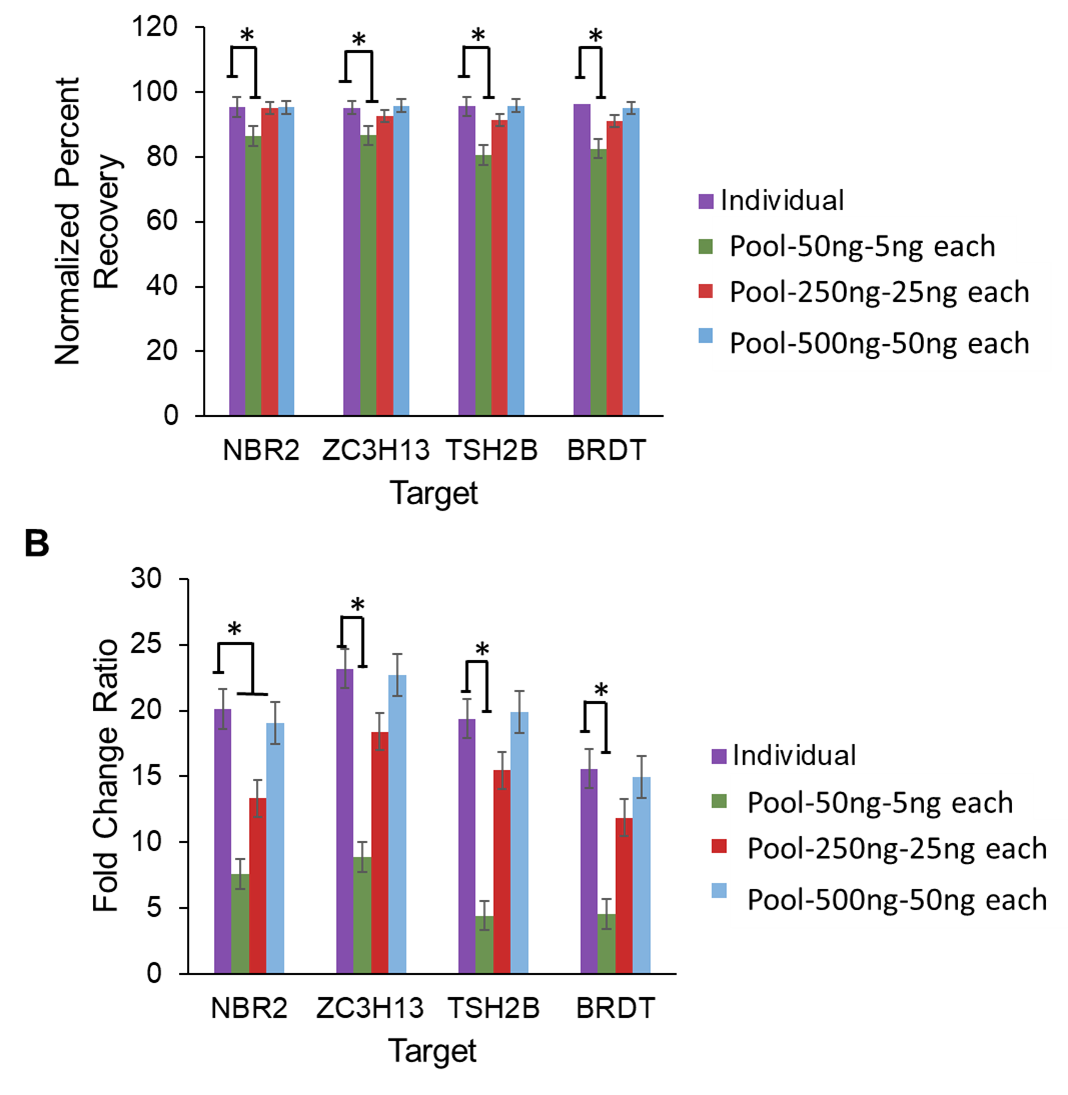

Figure S2. Effect of minimizing the amount of input DNA for Mx-MeDIP-Seq. Normalized percent recovery and fold change ratio for pools of DNA carried through Mx-MeDIP. Total amount of the pools of 50 ng, 250 ng, and 500 ng contained ten samples (5 ng, 25 ng, 50 ng, respectively). Statistical analysis between groups and individual were determined using one-way ANOVA and the Tukey test; p-values < 0.05 were considered statistically significant. For these studies, we carried out n=3 for each pool
